# Dimensionality Reduction of Diffusion MRI Measures for Improved Tractometry of the Human Brain

**DOI:** 10.1101/559351

**Authors:** Maxime Chamberland, Erika P. Raven, Sila Genc, Kate Duffy, Maxime Descoteaux, Greg D. Parker, Chantal M.W. Tax, Derek K. Jones

## Abstract

Various diffusion MRI measures have been proposed for characterising tissue microstructure over the last 15 years. Despite the growing number of experiments using different diffusion measures in assessments of white matter, there has been limited work on: 1) examining their covariance along specific pathways; and on 2) combining these different measures to study tissue microstructure. In this work, we first demonstrate redundancies in the amount of information captured by 10 diffusion tensor imaging (DTI) and high angular resolution diffusion imaging (HARDI) measures. Using a data-reduction approach, we identified two biologically-interpretable components that capture 80% of the variance in commonly-used DTI and HARDI measures profiled along 22 brain pathways extracted from typically developing children aged 8 - 18 years (n = 36). The first derived component captures properties related to hindrance and restriction in tissue microstructure, while the second component reflects characteristics related to tissue complexity and orientational dispersion. We demonstrate that the components generated by this approach preserve the biological relevance of the original measurements by showing age-related effects across developmentally sensitive pathways. Our results also suggest that HARDI measures are more sensitive at detecting age-related changes in tissue microstructure than DTI measures.

## 1. Introduction

The human brain is composed of multiple white matter fibres connecting grey matter areas dedicated to processes such as memory, cognition, language, or consciousness. Diffusion MRI (dMRI) (Basser et al., 1994, 2000; Basser and Jones, 2002; LeBihan et al., 2001) has become the prefered tool to probe the brain’s tissue microstructure non-invasively. Measures derived from diffusion tensor imaging (DTI) (Basser et al., 1994) can be obtained at each imaging voxel, including fractional anisotropy (FA) which reflects the degree of diffusion anisotropy (Pierpaoli and Basser, 1996), and mean diffusivity (MD), an indicator of the overall magnitude of diffusion. Based on local estimates of underlying trajectories at every voxel, dMRI is also capable of virtually reconstructing the structural architecture of the brain white matter pathways using tractography (Conturo et al., 1999; Mori and Van Zijl, 2002). The conventional approach to merge the quantitative nature of diffusion measures with the qualitative nature of tractography is to collapse voxel-based measures into a single scalar value per bundle (e.g., by averaging values over all vertices of a streamline; Jones et al. (2006); Kanaan et al. (2006); Jones et al. (2005a)). Individual differences in such summary diffusion-related measures can then be correlated, for example, with individual differences in cognition or behaviour. However, despite its well documented sensitivity, DTI has its limitations (Tournier et al., 2011; Jeurissen et al., 2013). For example, FA and MD lack specificity to the various physical properties of white matter, such as crossing fibres (Jeurissen et al., 2013), axon density and myelination (Beaulieu, 2002; Jones et al., 2013). Moreover, the average profile of those measures may vary along a given pathway depending on the underlying fibre architecture (Vos et al., 2012; Yeatman et al., 2012).

Recent advances in diffusion hardware, acquisition and modelling (Sotiropoulos et al., 2013; Jones et al., 2018; Assaf and Basser, 2005; Tournier et al., 2012; Jeurissen et al., 2014) have been introduced to overcome the limitations of DTI, giving access to previously inaccessible measures. High angular resolution diffusion imaging (HARDI; Tuch et al. (2002)) was originally developed to not only provide new anisotropy measures (Tournier et al., 2011) but also to solve the so-called *crossing fibre* problem, making tractography more robust (Descoteaux, 2015). Multi-shell acquisitions (Wedeen et al., 2005) have also facilitated new ways to link relevant tissue properties to the signal such as CHARMED (Assaf and Basser, 2005), AxCaliber (Assaf et al., 2008), ActiveAx (Alexander et al., 2010), multi-tensor models (Scherrer et al., 2016) and NODDI (Zhang et al., 2012) among others (for review, see Alexander et al. (2017)). In general, such models aim to extract parameters from intra- and extracellular compartments, and to estimate parameters such as axon diameter distributions and other high-order information.

Multi-shell acquisitions have also shown to improve the angular resolution of orientation distribution functions (ODFs) (Descoteaux et al., 2011; Jeurissen et al., 2014; Chamberland et al., 2018). In conjunction, new frameworks such as fixel-based analysis (Raffelt et al., 2012) have been proposed to map fibre-specific measures by looking at the apparent fibre density (AFD), a measure proportional to the underlying fibre density, as opposed to having voxel-specific scalar maps. The combination of frameworks such as along-tract profiling (Jones et al., 2005b; Corouge et al., 2006; Yeatman et al., 2012; De Santis et al., 2014; Colby et al., 2012; Cousineau et al., 2017) and tractometry (e.g., combining multiple measures (Bells et al., 2011)) allows for a comprehensive assessment of white matter microstructure. Both frameworks have the advantage of providing higher sensitivity to microstructural features of fibre pathways by mapping a set of MR-derived measures over white matter bundles. Recently, along-tract profiling has been successfully applied to study normal brain development (Geeraert et al., 2018) and to characterise areas of the brain with abnormal properties in various brain conditions (Dayan et al., 2016; Cousineau et al., 2017; Groeschel et al., 2014).

However, one problem arises with having access to multiple new measurements at each voxel and at each vertex forming a streamline: it quickly becomes intractable for existing analysis pipelines to process such high-dimensional data (a problem often refered to as the *curse of dimensionality*; Bellman (1961)), highlighting the need for new ways to visualise or analyse such data. Moreover, dMRI measures may share overlapping information which can cause redundancies in data analysis and ultimately decrease statistical power if strictly correcting for Type I errors (Penke et al., 2010; Metzler-Baddeley et al., 2017; Bourbon-Teles et al., 2017). A solution to this problem resides in dimensionality reduction, an established technique that has been successsfully applied in the past by the neuroimaging community (for review, see Mwangi et al. (2014)). Despite the growing number of experiments using different microstructural measures in assessments of white matter, there has been limited work on combining these different measures and on examining their covariance along specific pathways.

In this work, we explore the covariance of commonly-derived dMRI measures (De Santis et al., 2014). We propose a data reduction framework that takes advantage of those redundancies and aims to provide a better insight into patterns of associations between DTI and HARDI measures. Specifically, we identified common components that explain the maximal variance in measures profiled along multiple fibre bundles. We demonstrate the utility of our framework by showing enhanced sensitivity to the detection of age-related differences in tissue microstructure across developmentally sensitive pathways compared with the individual dMRI measures. Finally, we provide recommendations as to which set of measures are best suited for studies with limited capabilities in terms of data acquisition and processing.

## 2. Methods

### 2.1. Participants

This study reports on a sample of typically developing children aged 8 - 18 years (mean = 12.2 ± 2.8) participating in the Cardiff University Brain Research Imaging Centre (CUBRIC, School of Psychology) Kids study. The study was performed with ethics approval from the internal ethics review board and informed consent was provided from the primary caregiver of children enrolled in the study. Exclusion criteria included previous history of a neurological condition or epilepsy.

### 2.2. Data acquisition

Data from thirty-six (n = 36, 13 male) children were acquired using a multi-shell HARDI protocol on a Siemens 3T Connectom system with maximum gradient amplitude = 300 mT/m. The acquisition protocol consisted of 14 b_0_ images, 30 diffusion directions at b = 500, 1200 s/mm^2^ and 60 diffusion directions at b = 2400, 4000, 6000 s/mm^2^ with 2 × 2 × 2 mm^3^ voxels (TE/TR: 59/3000 ms, *δ*/∆: 7.0/23.3 ms).

### 2.3. Data pre-processing

Data quality assurance was performed on the raw diffusion volumes using slicewise outlier detection (SOLID; Sairanen et al. (2018)). Each dataset was then denoised in MRtrix (Veraart et al., 2016) and corrected for signal drift (Vos et al., 2017), subject motion (Andersson and Sotiropoulos, 2016), field distortion (Andersson et al., 2003), gradient non-linearities (Glasser et al., 2013; Suryanarayana et al., 2018) and Gibbs ringing artefacts (Kellner et al., 2016).

### 2.4. Local representation

Multi-shell multi-tissue constrained spherical deconvolution (MSMT-CSD; Jeurissen et al. (2014)) was applied to the pre-processed images to obtain voxel-wise estimates of fibre ODFs (fODFs; Tournier et al. (2004, 2007); Seunarine and Alexander (2009); Descoteaux et al. (2009)) with maximal spherical harmonics order *l*_*max*_ = 8. The fODFs were generated using a set of 3-tissue group-averaged response functions following image intensity normalisation in MRtrix (Tournier et al., 2012; Dhollander et al., 2016), enabling the direct comparison of fODF amplitudes across subjects. Diffusion tensors were also generated using linearly weighted least squares estimation (for b < 1200 s/mm^2^ data) providing the following quantitative scalar measures: FA, axial diffusivity (AD), radial diffusivity (RD), MD, geodesic anisotropy (GA; Fletcher et al. (2004)) and tensor mode representing the shape of the tensor (Kindlmann et al., 2007). In addition, HARDI measures were extracted from the fODFs of each subject. Those measures include fibre-specific AFD (Raffelt et al., 2012) for the bundles described in the next section, AFD_*tot*_ (spherical harmonics *l* = 0) and the Number of Fibre Orientations (NuFO) based on the number of local fODF peaks (Dell’Acqua et al., 2013). Finally, restricted signal fraction maps (FR, adapted from CHARMED to remove potential isotropic partial volume contamination; Assaf and Basser (2005)) were also computed using the fODFs peaks to initialise and regularise model-fitting. To summarise, ten dMRI measures related to tissue microstructure (*m* = 10) were generated for each subject.

### 2.5. Tractography and Tractometry

Whole-brain streamline tractography was perfomed using FiberNavigator (Chamberland et al., 2014) using 8 seeds/voxel evenly distributed across the whole brain (approximating 1.8M seeds), a minimum fODF amplitude of 0.1, a 1 mm step size (i.e. 0.5×voxel size), a 45° maximum curvature angle and streamlines whose lengths were outside a range of 20 mm to 300 mm were discarded. Twenty-two bundles of interest (*t* = 22) were then interactively dissected in the native space of each subject using a combination of include and exclude regions of interests (ROIs). Anatomical definitions and ROIs used to delineate each pathway are listed in the Supplementary Materials. The virtual dissection plan included:

**Commissural bundles**: anterior commissure (AC), body of the corpus callossum (CC), forceps minor (Genu), forceps major (Splenium).

**Association bundles** (bilateral): arcuate fasciculus (AF), cingulum (Cg), inferior fronto-occipital fasciculus (iFOF), inferior longitudinal fasciculus (ILF), optic radiations (OR), superior longitudinal fasciculus (SLF), uncinate fasciculus (UF).

**Projection bundles** (bilateral): corticospinal tract (CST), frontal aslant tract (FAT).

At this stage, we examined the covariance of the averaged diffusion measures for all bundles using Spearman’s correlation (r). Next, along-tract profiling was performed for each bundle using the Python toolbox developed by Cousineau et al. (2017), combined with DIPY functions (Garyfallidis et al., 2014). Bundles were first pruned to remove outliers as in (Cousineau et al., 2017; Garyfallidis et al., 2018). If necessary, the order in which the vertex-wise measures were stored was reversed, to ensure consistency across the subjects profiles (i.e., from left-to-right for commissural bundles, from inferior-to-superior for projection bundles and from posterior-to-anterior for association bundles). A representative core streamline was generated for each bundle (i.e., mean streamline of the pathway) and was subsequently resampled to *s* = 20 equidistant segments. Then, every vertex of every streamline forming the pathway was assigned to its closest segment along the core. The measure values of each vertex were then projected and averaged along each segment of the pathway, weighted by their geodesic distance from the core (Cousineau et al., 2017). An along-tract profile was finally generated for every combination of measure and pathway.

### 2.6. Dimensionality reduction

Each dataset comprised *m* = 10 dMRI-derived measures mapped along 440 white matter regions (*t* = 22 bundles × *s* = 20 segments). To explore the possible redundancy and complementarity of each measure, a principal component analysis (PCA) was performed on the concatenated set of profiles across subjects and bundles (Table 1) using the *tidy* data standard (Wickham et al., 2014). PCA reduces data dimensionality by extracting principal components that reflect relevant features in the data (Jolliffe, 2002; Abdi and Williams, 2010). The benefit is that a significant proportion of the variance in the data can be explained by a reduced number of orthogonal components, compared to the total number of raw input variables. PCA was performed by singular value decomposition of the z-transformed tract profiles via the *prcomp* package in R (RStudio Team, 2016). Here, the goal was to end up with the minimum number of components that summarise the maximum amount of information contained in the original set of diffusion measures. However, in order to avoid instability around the component loadings that comprise the principal components (Garg and Tai, 2013), measures showing significant covariance were discarded based on their correlation scores (r > 0.8) and the PCA was re-computed. Finally, the minimal number of principal components that accounted for the most variability was selected based on: 1) their interpretability (Metzler-Baddeley et al., 2017); and 2) the inspection of scree plots (Cattell, 1966) to select ranked components with an eigenvalue > 1.

**Table 1:** Data structure input for PCA. Individual subjects (n = 36), bundles (*t* = 22) and segments (*s* = 20) are concatenated to form observations while variables represent the measures (*m* = 10) derived from dMRI.

| Subject | Bundle | Section | FA | AD | ... | FR |
| --- | --- | --- | --- | --- | --- | --- |
| $S_1$ | Bundle<sub>1</sub> | Section<sub>1</sub> | FA<sub>111</sub> | AD<sub>111</sub> | ... | FR<sub>111</sub> |
| $S_2$ | Bundle<sub>1</sub> | Section<sub>1</sub> | FA<sub>211</sub> | AD<sub>211</sub> | ... | FR<sub>211</sub> |
| ⋮ | ⋮ | ⋮ | ⋮ | ⋮ | ... | ⋮ |
| $S_1$ | Bundle<sub>1</sub> | Section<sub>2</sub> | FA<sub>112</sub> | AD<sub>112</sub> | ... | FR<sub>112</sub> |
| ⋮ | ⋮ | ⋮ | ⋮ | ⋮ | ... | ⋮ |
| $S_n$ | Bundle<sub>b</sub> | Section<sub>s</sub> | FA<sub>nbs</sub> | AD<sub>nbs</sub> | ... | FR<sub>nbs</sub> |

### 2.7. Statistical analysis

PCA results were tested for sampling adequacy using a Kaiser-Meyer-Olkin (KMO; Dziuban and Shirkey (1974)) test followed by Bartlett’s test of sphericity to test whether the covariance matrix is significantly different from identity. We then ran an exploratory linear regression analysis to see whether profiles extracted from the PCA can provide increased sensitivity in the detection of age-related differences in tissue microstructure (as opposed to using the full set of *m* = 10 measures). It is important to recall that PCA results are always orthogonal, and therefore are statistically independent of one another. To address the multiple comparisons problem, a strict Bonferroni correction was applied to all linear models whereby statistical significance was defined as: p < 0.05 / (*m* measures × *t* bundles × *s* segments) resulting in p < 1.14e-5 for the ten raw measures, and p < 5.68e-5 for the first two principal components. All statistical analyses were carried out using RStudio v1.1.456 (RStudio Team, 2016).

## 3. Results

### 3.1. Measures covariance and profiling

The entire set of bundles and measures was successfully reconstructed in all subjects. Figure 1 shows the relationship between the various input measures averaged on different white matter pathways using a cross-correlation (Pearsons’s r) matrix representation. Matrices are re-organised using hierarchical clustering (Murtagh and Legendre, 2014), placing higher correlations closer to the diagonal in order to regroup measures that have similar correlations together, and push apart those that have the most dissimilar correlations. In general, the measures form two clusters across all bundles. The first cluster of positive correlations (r > 0.5) is observed between AD, FA, GA, AFD, AFD_*tot*_ and FR measures. The second cluster of positive correlations is formed of MD, RD and NuFO. The group-averaged diffusion measures of each bundle are reported in Suppl. Table 1.

**Figure 1:**
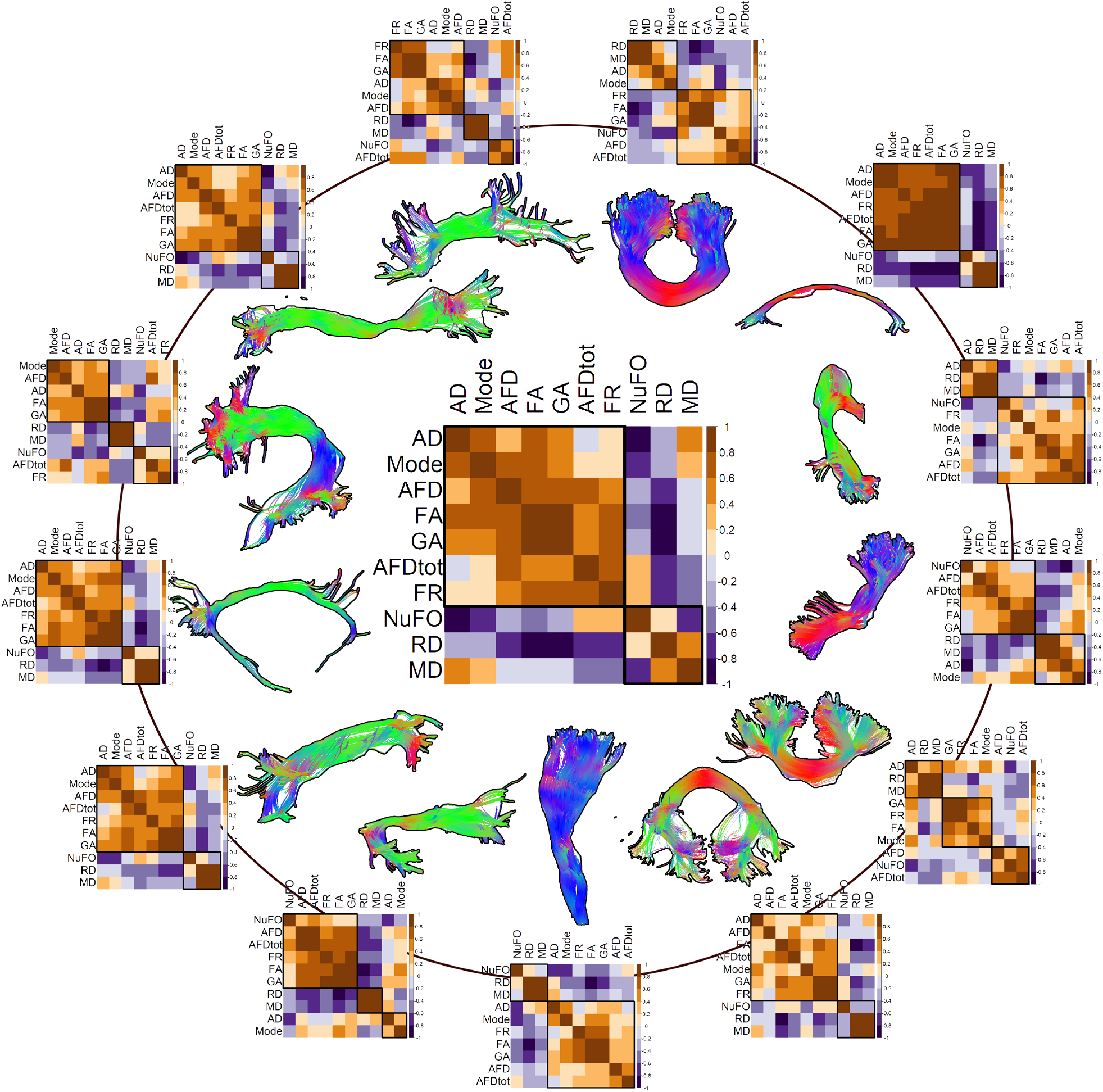
Correlation matrices of the ten diffusion measures, group-averaged for each extracted bundles. The middle image represents the average of all white matter bundles. Matrices are re-organised using hierarchical clustering, grouping measures that have similar correlations together. A first cluster of positive correlations (r > 0.5) is observed between most of the bundles for measures like AD, FA, GA, AFD, AFD_*tot*_, Mode and FR. A second set of positively correlated measures (NuFO, MD, RD) forms the second cluster. Note that for bilateral pathways, the left and right values were combined prior performing the correlation.

Important details about the spatial heterogeneity of the various input measures of interest appear when profiled along pathways. For example, FA, AFD and FR values all get progressively smaller along the CST as they approach the cortex (Figures 2, 3). Furthermore, the high number of fibre crossings near the centrum semiovale is reflected by a high NuFO index and is also marked by a low FA (Figure 3, black circles). As might be expected, HARDI-derived measures such as FR and AFD_*tot*_ seem to be less affected by the intra-voxel orientational heterogeneity of crossing regions than the tensor-based measures like FA, AD and RD. The correlation matrix in Figure 3 also highlights the similarity between the various microstructural profiles, indicating potential redundancies in the amount of information conveyed by those measures.

**Figure 2:**
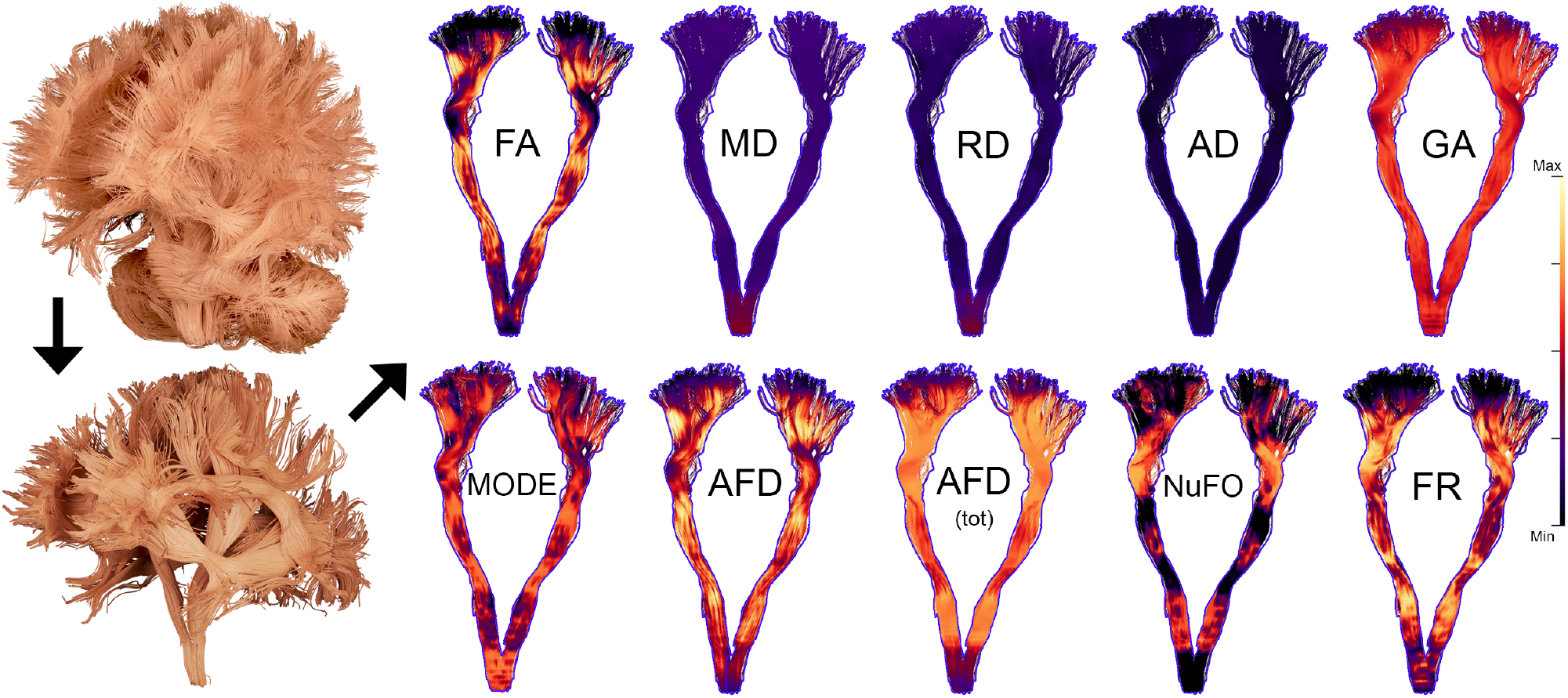
Overview of the ten input measures overlaid on the CST of a representative subject. Whole-brain tractograms (top-left) were manually dissected into *t* = 22 bundles (bottom-left) and measures were subsequently mapped along each pathway, providing information about their spatial heterogeneity.

**Figure 3:**
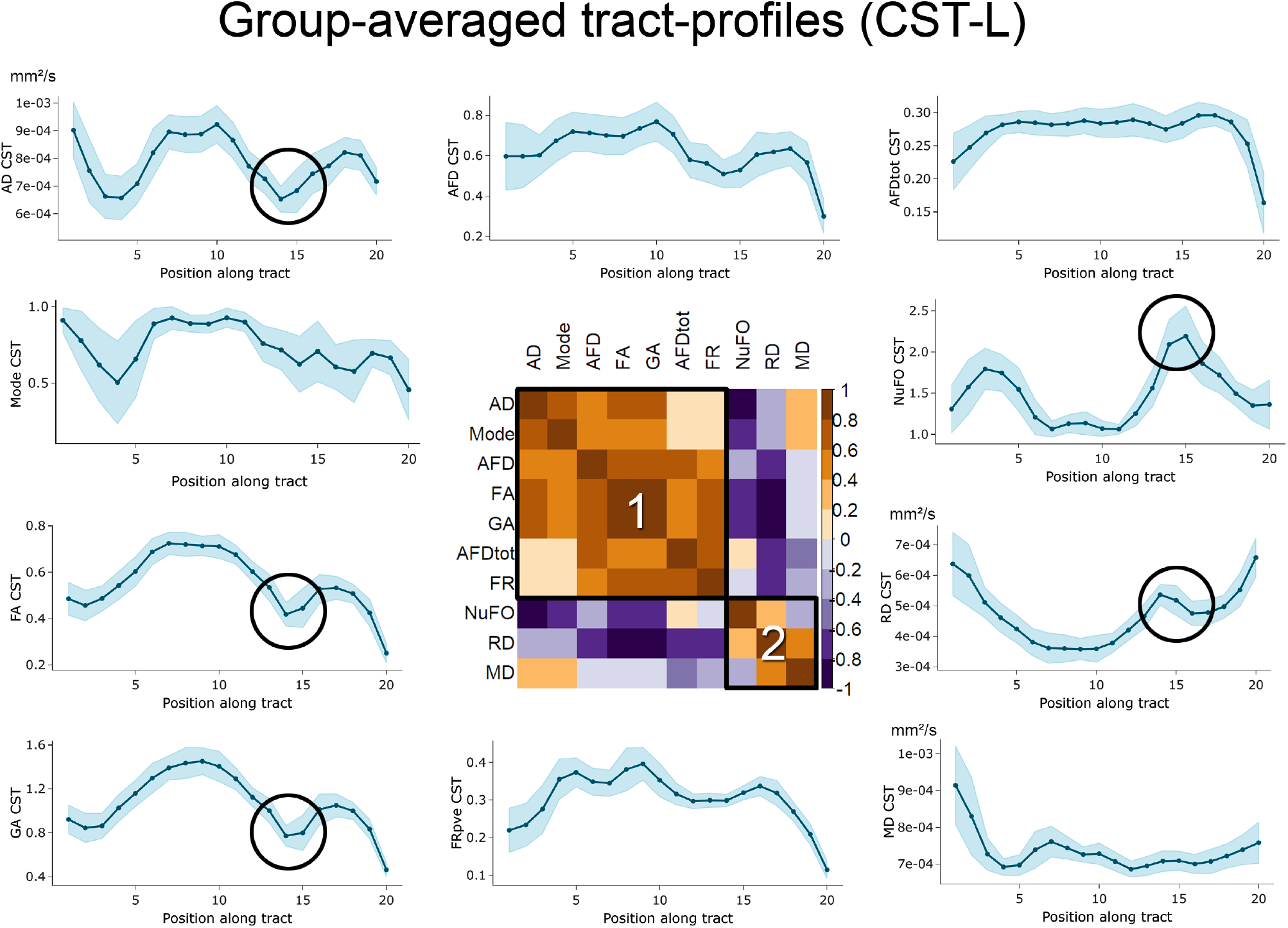
Group-average profiling of the ten input diffusion measures along the left CST for *s* = 20 segments, spanning from the brainstem (*s* = 1) to the cortex (*s* = 20). Heterogeneity in the profiles along the tract highlights the need for a vertex-wise assessment of the measures. Similarity between profiles also shows shared covariance between the measures, indicated by the two clusters (1 and 2) on the correlation matrix (sorting: hierarchical clustering). Shaded profile area: ±1 standard deviation.

### 3.2. Principal component analysis

The loading vector plots in Figure 4 show association patterns between the various input measures. The left panel shows PCA results performed on the entire set of measures. If two vectors subtend a small angle to each other, the two variables they represent are strongly correlated. When such vectors were found to be close, the one showing higher correlations with any other measures was removed. In line with the aforementioned results, shared covariance is observed between AD and tensor mode (r = 0.8), as well as between FA and GA (r= 0.95). After pruning up measures for multicollinearity (Figure 4, right panel), PCA results show that 80% of the variability in the data is accounted by the first two principal components (KMO: 0.64, sphericity: p < 2.2e-16). As shown in Figure 5, the PC that explains the largest proportion of the variance (PC1, 48%, *λ* = 3.4) is composed of hindrance-sensitive measures with AFD, FR and AFD_*tot*_ contributing positively (24%, 21% and 16%, respectively) and AD contributing negatively (25%). The second PC (PC2) represents 32% of the variance in the data (*λ* = 2.2) and is mostly driven by orientational dispersion and complexity-sensitive measurements, with its largest positive contribution from NuFO (34%), and negative contributions from AD (26%) and MD (25%) (Figure 5, PC2).

**Figure 4:**
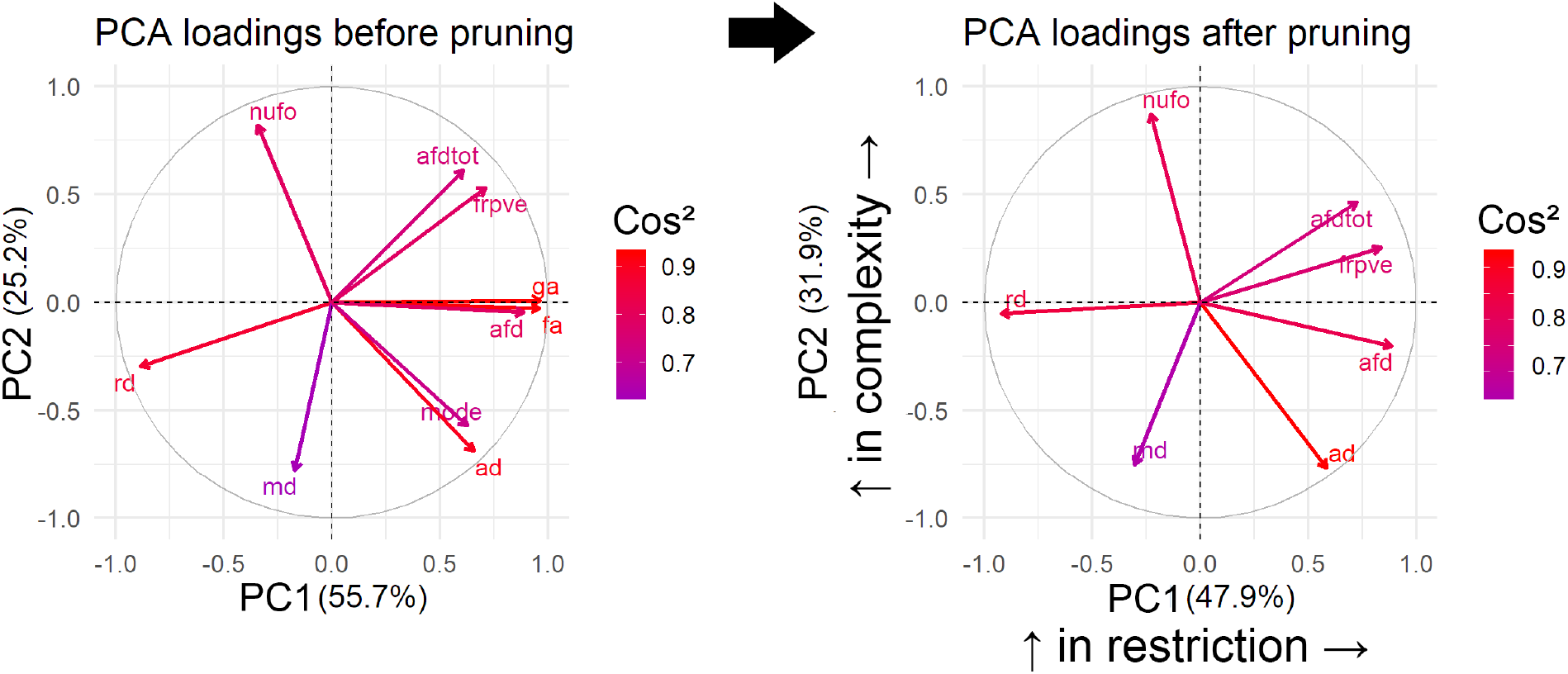
PCA results before (left) and after (right) multicollinearity analysis. To improve stability around PC1, AFD was kept over GA and FA due to its fibre specificity properties. Tensor mode was also discarded based on its collinearity with AD. On the right PCA, one can observe separation between the various measures, generating a hindrance-related component (PC1) that loads on AFD, AFD_*tot*_, RD and FR and a complexity-related component (PC2) that loads on NuFO, AD and MD. Here, the squared cosine notation (cos^2^) shows the importance of a measure for a given PC. A high cos^2^ indicates a good representation of the measure for a given principal component.

**Figure 5:**
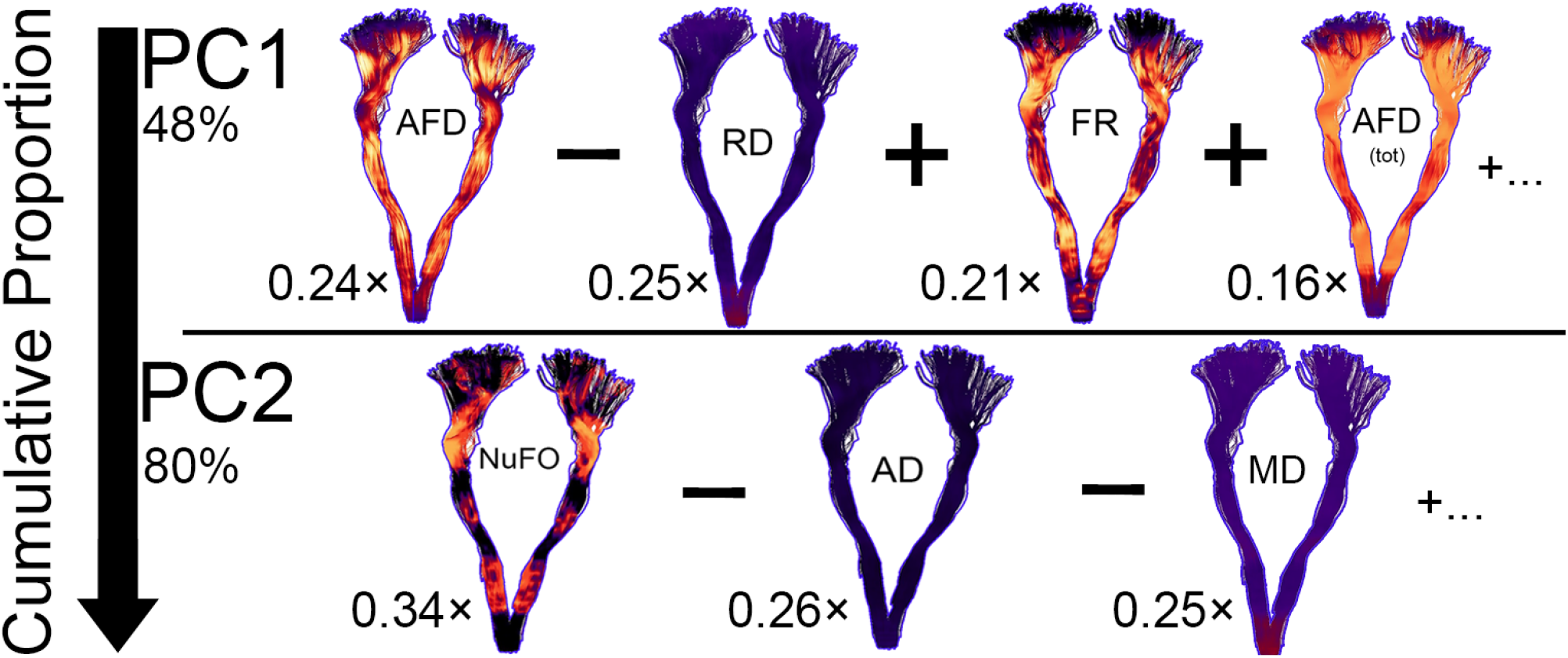
Visual overview of PC1 and PC2 with the contribution of each measure to that component. The first PC captures most of the hindrance- and restriction-related measures (AFD, RD, FR, AFD_*tot*_). The second PC mostly represents tissue complexity and orientational dispersion properties associated with NuFO, AD and MD. The first two components accounted for 80% of the variance in the diffusion measures.

### 3.3. Detecting differences in tissue microstructure

#### 3.3.1. Bundle averages

Here, we first report developmental changes in white matter tissue microstructure using bundle-averaged measures (*m* = 10, *s* = 1) and PCA components (*m* = 2, *s* = 1). Significance thresholds were Bonferroni-corrected to account for multiple comparisons (p < 2.27e-4 for the ten raw measures, and p < 1.14e-3 for the two principal components). Figure 6 shows a significant increase in PC1 as a function of age for the left iFOF and CST, whereas no correlation with age was observed between individual hindrance-related measures. Significant positive correlations were found between PC1 (restriction-related component) and age in the following language-related pathways: AF (left: R^2^: 0.34, p = 1.06e-4), iFOF (left: R^2^: 0.31, p = 2.51e-4), FAT (left: R^2^: 0.43, p = 9.06e-6, right: R^2^: 0.43, p = 7.77e-6), UF (right: R^2^: 0.26, p = 9.76e-4) and motor pathways: CST (left: R^2^: 0.40, p = 1.92e-5, right: R^2^: 0.40, p = 2.0e-5), CC (R^2^: 0.29, p = 4.51e-4). One significant positive correlation between PC2 (dispersion-related component) and age was found in the SLF (right: R^2^: 0.27, p = 6.82e-4).

**Figure 6:**
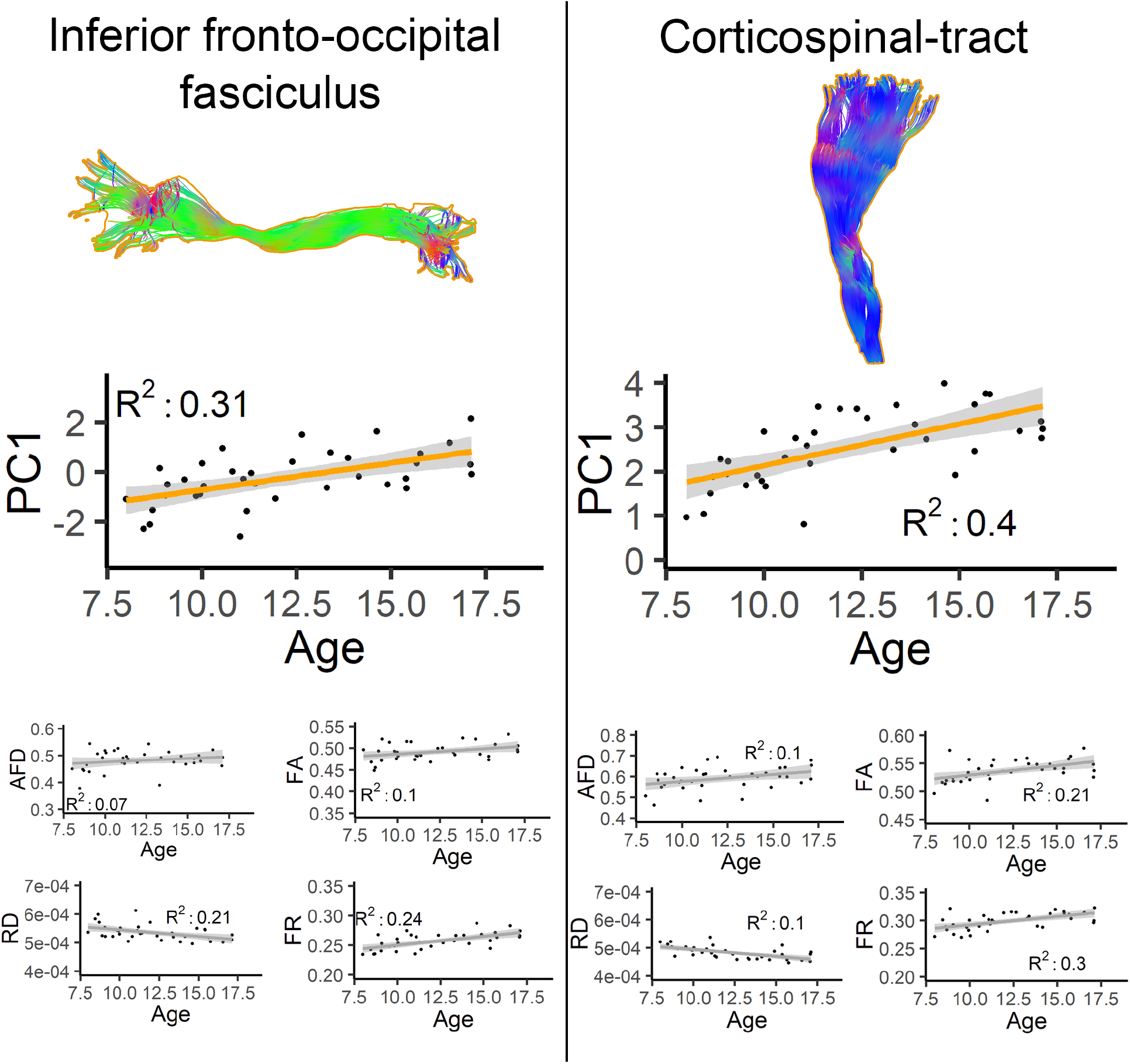
Developmental changes in iFOF and CST bundles. PC1 show significant positive correlation with age, whereas no correlation was observed between the individual hindrance-sensitive measures.

**Figure 7:**
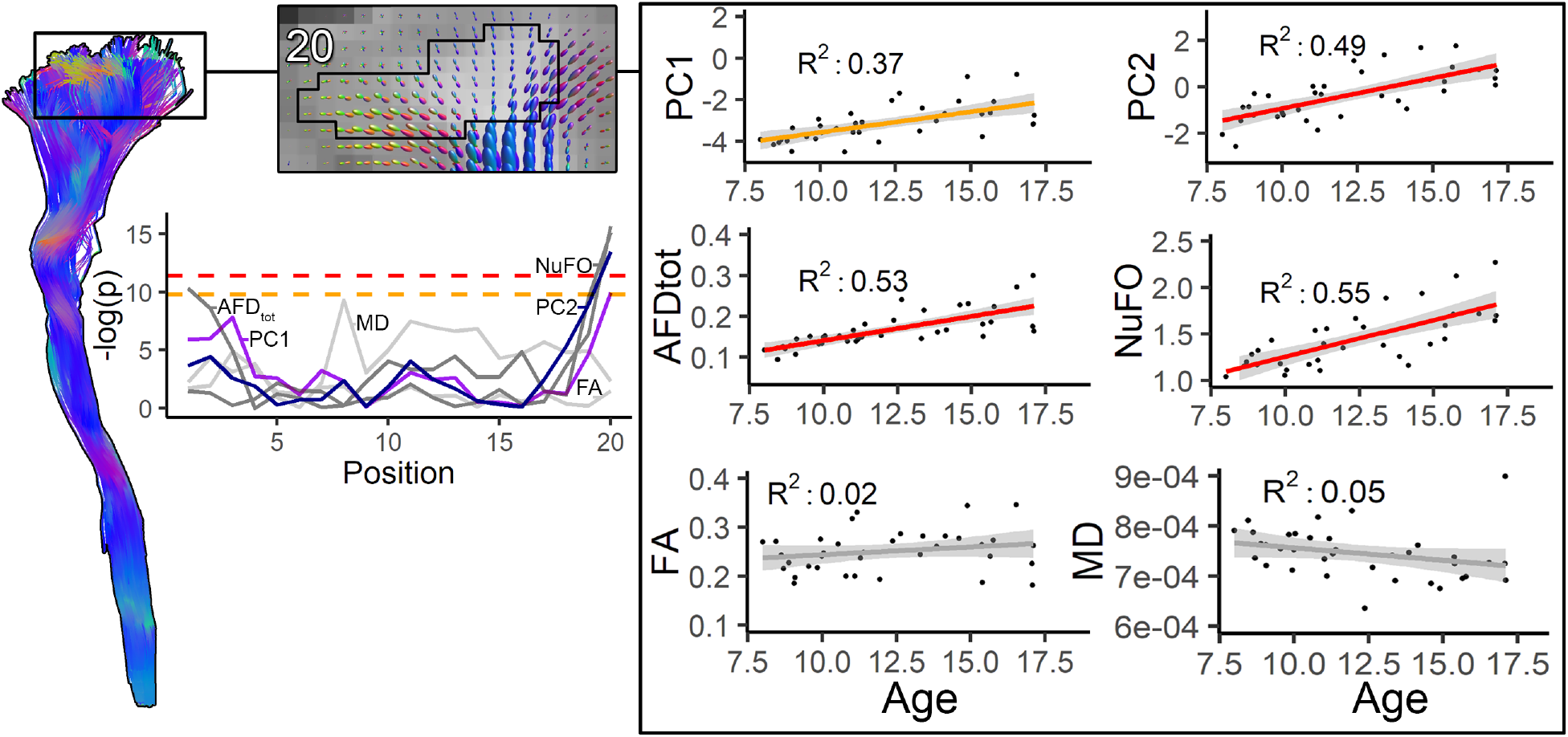
Age relationships captured by PC1 and PC2 over the left CST. Highlighted section of an axial slice overlaid with fODFs reconstruction of a representative participant (top left). Contoured area shows where streamlines terminated to form segment 20. At the group level, significant positive correlations with age were found with PC1 and PC2 (top right). Significant positive correlations were also found for HARDI measures AFD_*tot*_ and NuFO (middle right). No significant correlations were observed for any of the DTI measures (bottom right). Profile plots indicate where significant differences in tissue microstructure were located along the CST (-log(p) scale).

No significant age relationships were found between any of the bundles and FA, GA, Mode, AFD, AFD_*tot*_, FR and NuFO. Significant negative correlations with age were found in RD for the AF (left: R^2^: 0.37, p = 5.40e-5, right: R^2^: 0.34, p = 1.02e-4), Cg (left: R^2^: 0.33, p = 1.45e-4, right: R^2^: 0.47, p = 2.22e-6), CST (left: R^2^: 0.35, p = 7.36e-5, right: R^2^: 0.33, p = 1.28e-4), FAT (left: R^2^: 0.37, p = 4.65e-5, right: R^2^: 0.28, p = 4.67e-4). Significant negative correlations with age were found in MD for the FAT (left: R^2^: 0.33, p = 1.22e-4, right: R^2^: 0.31, p = 2.18e-4), AF (left: R^2^: 0.39, p = 2.48e-5, right: R^2^: 0.36, p = 6.28e-5), CST (left: R^2^: 0.40, p = 2.06e-5, right: R^2^: 0.42, p = 1.12e-5), SLF (right: R^2^: 0.38, p = 3.21e-5) and Cg (right: R^2^: 0.37, p = 4.06e-5). One significant negative correlation was found between AD and age in the SLF (right: R^2^: 0.41, p = 1.80e-5).

#### 3.3.2. Along-tract profiling

Here, we report on developmental changes in tissue microstructure seen with along-tract profiling. Table 2 reports the measures and tract segment mapped along different pathways where significant correlation with age was observed. Significant positive correlations were found between PC1 (*restricted* component) and age near the motor cortex area for the CST_20_ (right: R^2^: 0.37, p = 4.91e-5) and CC_3_ (R^2^: 0.37, p = 4.62e-5). Significant age-related positive correlations with PC2 were observed for motor-related pathways: CC_2_ (R^2^: 0.37, p = 5.63e-5), CST_20_ (right: R^2^: 0.49, p = 1.41e-6), CST_19_ (left: R^2^: 0.43, p = 8.59e-6) and language-related pathways: FAT_19_ (right: R^2^: 0.38, p = 4.08e-5), iFOF_19_ (right: R^2^: 0.39, p = 2.89e-5), ILF_1_ (right: R^2^: 0.35, p = 9.26e-5), UF_3,4_ (right: R^2^: 0.43 & 0.39, p = 7.71e-6 & 3.0e-5) and SLF_1,2_ (R^2^: 0.46 & 0.42, p = 3.41e-6 & 1.14e-5).

**Table 2:** Segments of white matter bundles where a significant correlation between diffusion measures and age were observed. Subscript ordering for along-tract positions: left (*s* = 1) to right (*s* = 20) for commissural bundles, inferior (*s* = 1) to superior (*s* = 20) for projections bundles and posterior (*s* = 1) to anterior (*s* = 20) for associations bundles. Positive / negative correlations are indicated by an increasing /decreasing arrow, respectively. Significance thresholds for the measures and components were set as p < 1.13e-5 and p < 5.68e-5, respectively (adjusted R^2^ > 0.3).

| Individual diffusion measures |  |  |  |  | PCA |  |
| --- | --- | --- | --- | --- | --- | --- |
| RD | MD | AFD<sub>tot</sub> | NuFO | FR | PC1 | PC2 |
| ↘ r-FAT<sub>8</sub> | ↘ r-AF<sub>16</sub> | ↗ l-CST<sub>1,20</sub> | ↗ CC<sub>1</sub> | ↗ r-FAT<sub>6</sub> | ↗ CC<sub>3</sub> | ↗ CC<sub>2</sub> |
|  | ↘ l-Cg<sub>7</sub> | ↗ r-CST<sub>20</sub> | ↗ r-CST<sub>20</sub> |  | ↗ r-CST<sub>20</sub> | ↗ r-CST<sub>20</sub> |
|  |  |  |  |  |  | ↗ l-CST<sub>19</sub> |
|  |  |  |  |  |  | ↗ r-FAT<sub>19</sub> |
|  |  |  |  |  |  | ↗ r-IFOF<sub>19</sub> |
|  |  |  |  |  |  | ↗ r-ILF<sub>1</sub> |
|  |  |  |  |  |  | ↗ r-UF<sub>3,4</sub> |
|  |  |  |  |  |  | ↗ r-SLF<sub>1,2</sub> |

No significant age relationships were found in any bundles for FA, GA, Mode, AD and AFD. Significant negative correlations were observed for RD in the FAT_8_ (right: R^2^: 0.42, p = 1.08e-5) and for MD in the AF_16_ (right: R^2^: 0.44, p = 5.62e-6) and Cg_7_ (left: R^2^: 0.43, p = 6.5e-6). Significant positive correlations were also found for AFD_*tot*_ in the CST_20_ (left: R^2^: 0.50, p = 9.02e-7, right: R^2^: 0.53, p = 2.73e-7) and CST_1_ (left: R^2^: 0.46, p = 2.88e-6). For NuFO, significant age-related differences in tissue complexity were observed in the CST_20_ (right: R^2^: 0.54, p = 1.56e-7) and CC_3_ (R^2^: 0.43, p = 7.09e-6). Finally, one significant positive correlation in FR was found for the FAT_6_ (right: R^2^: 0.57, p = 4.9e-8).

## 4. Discussion

### 4.1. Extraction of interpretable components

The aim of this study was to systematically examine any potential covariance between various diffusion measures mapped along white matter fibre bundles extracted from a cohort of typically-developing children and adolescents. We first examined the covariance of the measures averaged over different bundles, which revealed two clusters of interdependent measures. The first cluster revealed that measures sensitive to restricted diffusion shared high correlations with each other. Similarly, measures which are known to be sensitive to local complexity or orientational dispersion co-varied. When profiled along pathways, measures showed heterogeneity across the trajectory of diverse pathways, but their interdependency remained marked by a two-cluster formation. This provided motivation for the next step of our analysis, where we performed a PCA to collapse the inter-dependent measures into the principal modes of variance. This was done by profiling multiple brain fibre systems based on their dMRI features, and then deriving a set of principal components that best represent those individual measures. We then showed the sensitivity of these new components to the detection of differences in tissue microstructure of white matter pathways by exploring their relationship with the age of participants.

A common problem with PCA is that the interpretation of the resulting components can be challenging. Here, the principal components loaded onto variables that shared similarities in their sensitivity to different tissue properties, making the interpretation of the resulting components more meaningful. Measures accounting for the largest percentage of variance in the data (forming PC1) are those known to be most sensitive to hindrance or restriction in the signal, including RD, AFD and FR. In contrast, PC2 features measures that could reflect complexity or orientational dispersion in the signal, such as NuFO, AD and MD.

The raw tract-averaged diffusion measures showed significant negative correlations in MD and RD with age across a range of developmentally sensitive tracts, which is in line with previous studies that also report a decrease in MD and RD with age, whereas FA shows slower increases with age in late childhood (Lebel et al., 2008). Additionally, we observed significant age relationships for PC1 and PC2 in developmental-sensitive tracts related to language (i.e., AF, SLF, FAT, iFOF, ILF) and motor functions (i.e., CST and CC), which is also in line with previous reports in the literature (Genc et al., 2017, 2018; Lebel et al., 2017; Geeraert et al., 2018; Genc et al., 2018). We note that sex differences were not accounted for due to our relatively small sample size but may play a role in some of the differences we observed across pathways (Seunarine et al., 2016). Our results highlight the sensitivity of PC1 and PC2 as a composite measure by 1) showing significant correlation with age in regions where other measures did not, and 2) reflecting effects captured by the other measures. Examining the tract-averaged values for PC2, only one significant correlation with age was found for the right SLF. However, additional significant correlation between age and PC2 were observed when performing along-tract profiling. A potential explanation for this findings resides in the nature of PC2, with most of its contributions coming from AD, MD and NuFO. Indeed, since PC2 reflects the local complexity at each voxel, measures like NuFO will vary depending on the underlying structural architecture and therefore taking the average value across the bundle may lead to a summary statistic that is hard to interpret. In contrast, the values of PC1 (or AFD) remain relatively constant over bundles and thus are less impacted by calculating the bundle average. This may also explain why the first principal component derived from bundle-averages was found to show significant correlations with age in white matter bundles, whereas the original DTI measures did not.

Restricted (or hindered) diffusion is primarily caused by dense packing of axons and their cell membranes (Beaulieu, 2002). Other tissue properties such as myelination and local complexity can also affect the degree of hindrance or restrictance measured at each voxel (Vos et al., 2012). In the current study, an increase in PC1 may indicate higher coherence of the underlying white matter bundles for our older subjects, in comparison with the younger ones. Previous studies have demonstrated that dMRI measures can be sensitive to age-related differences, and those are often associated with an increased microstructural organisation (for review, see Lebel et al. (2017)). Given the well-established role of the CST in supporting motor performance, our finding of increased hindrance with age in typically developing children is in line with previous research that showed that brain maturation varies across different pathways, with commissural and projection tracts reaching maturation by early adolescence while association pathways develop over a longer time period (Geeraert et al., 2018). Interestingly, PC2 also captured an increase in orientational dispersion for that same region (which was either marked by an increase in NuFO or decrease in MD, Figure 4). The fact that those changes appear near the cortex, a region usually contaminated by partial volume effects, highlights the role of MSMT-CSD in achieving adequate fODFs representation at the boundary between gray matter and white matter (Jeurissen et al., 2014).

### 4.2. Choice of measures

A growing interest in utilising advanced dMRI measures to study the human brain motivated us to investigate the shared relationship between DTI and HARDI measures (De Santis et al., 2014). Being a relatively fast-developing field, dMRI offers a multitude of mathematical models to represent the underlying tissue microstructure (Alexander et al., 2017). Here, we focused on DTI and HARDI measures (Descoteaux, 2015). dMRI measures are generally sensitive to differences in tissue microstructure that can potentially be linked to fibre properties such as myelination and axon density (Scholz et al., 2014). Despite the fact that the specific interpretation of these measures remains controversial (Jones et al., 2013), DTI and HARDI measures are routinely used by neuroscientists and clinicians to gain insights into white matter properties. The findings reported here are in line with existing evidence suggesting that HARDI measures may be more specific than DTI for the detection of differences in tissue microstructure (Tournier et al., 2011; Jeurissen et al., 2013; Cousineau et al., 2017). Our results suggest that combining the sensitivity of DTI and the specificity of HARDI has the potential for compromise between the two techniques. Other macrostructural measures (e.g., bundle volume or mean length; Lebel et al. (2012, 2008, 2017); Geeraert et al. (2018); Girard et al. (2014)) have been used to study brain development and may also provide complementary features that could be applied in the proposed framework. Other measures such as rotationally invariant spherical harmonic features (RISH; Mirzaalian et al. (2015)) could also be introduced in the current framework, with the main advantage of representing more directly the diffusion signals rather than relying on various microstructural models.

Ultimately, the key challenge resides in knowing what measure (or combination of; De Santis et al. (2014)) provides the best value in terms of scanning and processing time. In other words, one may ask: which set of microstructural measures provides the most variance per unit of scan time? Indeed, optimising scan time by knowing in advance which measures will provide better sensitivity to the current study is challenging, since this information cannot be determined *a priori*. One approach (although not practical) would be to acquire all possible data and fit all existing models using a brute force approach, followed by extensive exploratory analyses. A better suited alternative resides in the careful design of dMRI studies. To help with the planning of future studies and based on our observations, we present some recommendations for data analysis.

#### DTI vs HARDI

DTI measures can nowadays be easily be derived from a conventional 30 directions protocol acquired at b = 1000 s/mm^2^ in approximatively five minutes. That being said, instead of relying on DTI-based measures (e.g., FA and MD), a hindrance-sensitive measure such as AFD (Raffelt et al., 2012; Dell’Acqua et al., 2013) and a complexity-sensitive one such as NuFO (Dell’Acqua et al., 2013) are both adequate and accessible alternatives. Indeed, CSD can usually be performed on data with a minimum of 45-60 directions acquired at b = 1000 s/mm^2^ (Dell’Acqua and Tournier, 2018; Alexander and Barker, 2005). Moreover, going beyond single b-value acquisitions will provide a better estimation of partial volume effects and better characterisation of various tissue types that will subsequently improve HARDI reconstructions in those areas (Jeurissen et al., 2014; Chamberland et al., 2018).

#### Tract-average vs along-tract profiling

In the context of along-tract profiling, age-related effects might be more pronounced when performing group-wise comparisons (e.g., young vs old, patients vs controls; Yeatman et al. (2014)) rather than directly looking at a single cross-sectional change in tissue microstructure. Indeed, theses changes might be too subtle to detect, especially considering that the age range of our participants falls on the *inflection point* of the developmental curve (Lebel and Beaulieu, 2011). Moreover, one may consider first looking at the profile along each tract of interest and ask the following: are there any benefits in sub-segmenting the profile into finer portions? Admittedly, if the measure of interest remains stable along the pathways, a conventional tract average is probably better suited than looking at a constant profile at multiple points. Concomitantly and depending on the research hypothesis, the use of a more permissive approach such as false discovery rate (FDR) correction should be considered to assess differences along multiple adjacent bundle segments.

### 4.3. Future perspectives

Over the last few years, a wide range of supervised and unsupervised learning applications based on feature extraction has emerged, ranging from individual classifiers for specific brain disorders (Wang et al., 2010; Chu et al., 2012) to data predictors of brain function (Chen et al., 2009; Franke et al., 2012; Casanova et al., 2011; Kucukboyaci et al., 2014). In the field of functional MRI, independent component analysis (ICA) is a successful example of unsupervised dimensionality reduction that allows the extraction of temporally segregated resting-state networks (Beckmann et al., 2009). In all cases, data reduction approaches facilitate a stream to analyse and interpret the increasingly large multi-dimensional data generated by new methodological models. Admittedly, despite PCA being one of the most commonly-used tool for data reduction, it can also over-fit data (Kramer, 1991) and potentially require multiple post-processing regression steps to explore the link between the resulting components and the observations. Similar to PCA, a canonical correlation analysis (Hotelling, 1936) may help in finding the link between the correlated measures and observations by extracting their joint information. Other non-linear dimensionality reduction techniques based on manifold learning such as isometric feature mapping (Tenenbaum, 1998) or locally linear embedding (Roweis and Saul, 2000) may also better disentangle the measurement space. However, one has to be cautious when applying advanced models of dimensionality reduction to medical imaging as it is often a trade-off between model accuracy and interpretability. Indeed, although these techniques may result in better disentangling of the manifold space, this often comes at the expense of generating complex and less interpretable features that cannot be related to brain tissue microstructure.

Overall, data representation frameworks such as the one presented here can become fundamental in advancing the application of diffusion models in health and disease. The proposed framework may open new avenues for examining brain microstructure in general and other related lines of research, especially if complimented with other modalities such as measures derived from quantitative magnetisation transfer (Rovaris et al., 2003; Cercignani and Bouyagoub, 2018). Indeed, with the ever-growing acquisition of large cohorts of subjects, feature extraction techniques may become essential tools for processing multi-dimensional brain imaging datasets (e.g., the Human Connectome Project with >1,000 young adults scans; Van Essen et al. (2013) or the UK Biobank with its 500,000 participants; Miller et al. (2016)).

Finally, our study may also open new avenues for fibre clustering by leveraging microstructural properties mapped over fibre bundles. Suppl. Figure 1 shows how different bundles project and cluster in the new reference frame formed by PC1 and PC2. One can observe that PC1 is sensitive to various hindrance level in white matter by disentangling bundles such as the CST (green), genu (blue) and splenium (pink). Conversely, pathways that are known to have many crossing regions such as the AF and the SLF are located on the superior portion of the bi-plot, showing properties of increased orientational dispersion (Suppl. Figure 1 orange and purple, respectively).

## 5. Conclusions

In summary, our findings demonstrate that there exists redundancies in measures conventionally derived from dMRI and that those redundancies may be exploited to reduce the risk of Type I errors, arising from multiple statistical comparisons. Our results support the use of data reduction to detect along-tract differences in tissue microstructure. Specifically, the curse of dimensionality and redundancies in statistical analyses were considerably mitigated by extracting components sensitive to diffusion properties of hindrance and complexity. From an application perspective, a general increase in white matter hindrance was found to have a significant correlation with age in various developmentally sensitive pathways, a change that would otherwise remain undetected using conventional approaches. Under limited acquisition or processing capabilities, our results indicate that one could invest in deriving a hindrance-sensitive HARDI measure (e.g., AFD or FR) and a complexity-, or dispersion-sensitive measure (e.g., NuFO or MD) to study tissue-microstructure.

## Acknowledgements

The authors would like to thank Umesh Rudrapatna, Peter Hobden, John Evans, Alison Cooper and Isobel Ward (CUBRIC) for their support with data acquisition. The authors are also thankful to Jean-Christophe Houde (Sherbrooke Connectivity Imaging Lab) for sharing implementation details. The authors are also grateful to the Natural Sciences and Engineering Research Council of Canada (NSERC), the Marshall Sherfield Postdoctoral Fellowship (UK), the Netherlands Organisation for Scientific Research (NWO), the Wellcome Trust (UK) and the Engineering and Physical Sciences Research Council (EPSRC, UK) for supporting this research. Finally, we thank the participants and their families for participating in the study.

## Supplementary Data

### Supplementary Table 1

**Table S1:** Quantitative overview of the group-averaged dMRI measures across all bundles (mean±std). AD, RD and MD are expressed in mm^2^/s ×10^*-*4^.

| Bundle | FA | AD | RD | MD | GA | Mode | AFD | AFD<sub>tot</sub> | NuFO | FR |
| --- | --- | --- | --- | --- | --- | --- | --- | --- | --- | --- |
| AC | 0.45±0.14 | 8.1±1.2 | 5.9±1.3 | 8.0±0.9 | 0.82±0.28 | 0.68±0.43 | 0.44±0.18 | 0.21±0.05 | 1.4±0.5 | 0.21±0.08 |
| AF (l) | 0.47±0.13 | 7.5±1.4 | 5.3±0.9 | 7.3±0.4 | 0.90±0.28 | 0.51±0.50 | 0.46±0.20 | 0.25±0.05 | 1.7±0.7 | 0.29±0.09 |
| AF (r) | 0.46±0.14 | 7.3±1.5 | 5.2±0.9 | 7.1±0.4 | 0.86±0.29 | 0.49±0.52 | 0.44±0.20 | 0.25±0.05 | 1.8±0.7 | 0.28±0.09 |
| CC | 0.53±0.19 | 8.7±2.0 | 5.0±1.6 | 7.7±1.2 | 1.06±0.44 | 0.68±0.44 | 0.55±0.21 | 0.25±0.06 | 1.4±0.6 | 0.29±0.12 |
| CST (l) | 0.54±0.17 | 7.7±1.5 | 4.8±1.5 | 7.4±1.1 | 1.02±0.37 | 0.70±0.41 | 0.59±0.21 | 0.27±0.05 | 1.5±0.7 | 0.30±0.10 |
| CST (r) | 0.53±0.17 | 7.6±1.6 | 4.8±1.4 | 7.3±1.0 | 1.01±0.37 | 0.69±0.42 | 0.58±0.20 | 0.26±0.05 | 1.6±0.7 | 0.30±0.10 |
| ILF (l) | 0.47±0.13 | 8.1±1.4 | 5.6±0.9 | 7.8±0.6 | 0.89±0.28 | 0.57±0.47 | 0.42±0.18 | 0.23±0.05 | 1.5±0.6 | 0.24±0.09 |
| ILF (r) | 0.47±0.13 | 7.9±1.3 | 5.5±0.9 | 7.6±0.6 | 0.88±0.28 | 0.54±0.49 | 0.41±0.18 | 0.23±0.06 | 1.5±0.6 | 0.25±0.09 |
| OR (l) | 0.49±0.15 | 8.4±1.9 | 5.3±1.1 | 7.7±0.8 | 0.92±0.33 | 0.65±0.47 | 0.46±0.21 | 0.23±0.06 | 1.4±0.6 | 0.25±0.10 |
| OR (r) | 0.49±0.15 | 8.1±1.7 | 5.3±1.1 | 7.6±0.8 | 0.92±0.33 | 0.63±0.46 | 0.45±0.21 | 0.24±0.06 | 1.5±0.6 | 0.26±0.10 |
| SLF (l) | 0.42±0.14 | 7.2±1.3 | 5.5±0.9 | 7.2±0.5 | 0.81±0.28 | 0.43±0.52 | 0.38±0.18 | 0.24±0.07 | 1.8±0.7 | 0.26±0.09 |
| SLF (r) | 0.44±0.14 | 7.3±1.4 | 5.3±0.9 | 7.2±0.5 | 0.85±0.29 | 0.46±0.51 | 0.40±0.19 | 0.24±0.06 | 1.7±0.7 | 0.27±0.09 |
| UF (l) | 0.44±0.12 | 8.3±1.2 | 5.9±0.9 | 8.0±0.5 | 0.77±0.24 | 0.67±0.40 | 0.40±0.18 | 0.21±0.06 | 1.3±0.5 | 0.19±0.07 |
| UF (r) | 0.46±0.12 | 8.2±1.3 | 5.6±0.9 | 7.7±0.5 | 0.79±0.24 | 0.72±0.37 | 0.44±0.18 | 0.22±0.05 | 1.3±0.6 | 0.19±0.06 |
| FAT (l) | 0.44±0.15 | 7.6±1.4 | 5.5±1.0 | 7.4±0.5 | 0.82±0.31 | 0.52±0.48 | 0.41±0.20 | 0.23±0.07 | 1.6±0.7 | 0.25±0.10 |
| FAT (r) | 0.43±0.14 | 7.5±1.4 | 5.5±1.0 | 7.3±0.5 | 0.81±0.30 | 0.51±0.49 | 0.42±0.20 | 0.24±0.07 | 1.7±0.7 | 0.25±0.09 |
| Cg (l) | 0.48±0.15 | 8.3±1.5 | 5.4±1.0 | 7.7±0.5 | 0.91±0.32 | 0.67±0.44 | 0.47±0.20 | 0.22±0.05 | 1.5±0.6 | 0.26±0.09 |
| Cg (r) | 0.46±0.13 | 8.1±1.3 | 5.5±0.9 | 7.7±0.5 | 0.85±0.28 | 0.63±0.46 | 0.44±0.18 | 0.21±0.05 | 1.5±0.6 | 0.24±0.08 |
| Genu | 0.55±0.19 | 8.5±1.7 | 4.9±1.6 | 7.6±0.8 | 1.03±0.44 | 0.68±0.46 | 0.48±0.20 | 0.23±0.05 | 1.4±0.6 | 0.27±0.11 |
| iFOF (l) | 0.49±0.14 | 8.5±1.6 | 5.4±1.0 | 7.7±0.6 | 0.93±0.30 | 0.67±0.42 | 0.48±0.19 | 0.24±0.05 | 1.5±0.6 | 0.26±0.09 |
| iFOF (r) | 0.50±0.14 | 8.3±1.5 | 5.2±1.0 | 7.6±0.6 | 0.93±0.31 | 0.65±0.44 | 0.48±0.20 | 0.24±0.05 | 1.5±0.6 | 0.26±0.09 |
| Splenium | 0.61±0.21 | 9.6±2.1 | 4.5±2.0 | 7.8±1.4 | 1.25±0.52 | 0.78±0.39 | 0.56±0.22 | 0.25±0.06 | 1.2±0.5 | 0.32±0.13 |

### Supplementary Figure1

**Figure 1: Supplementary Figure 1:**
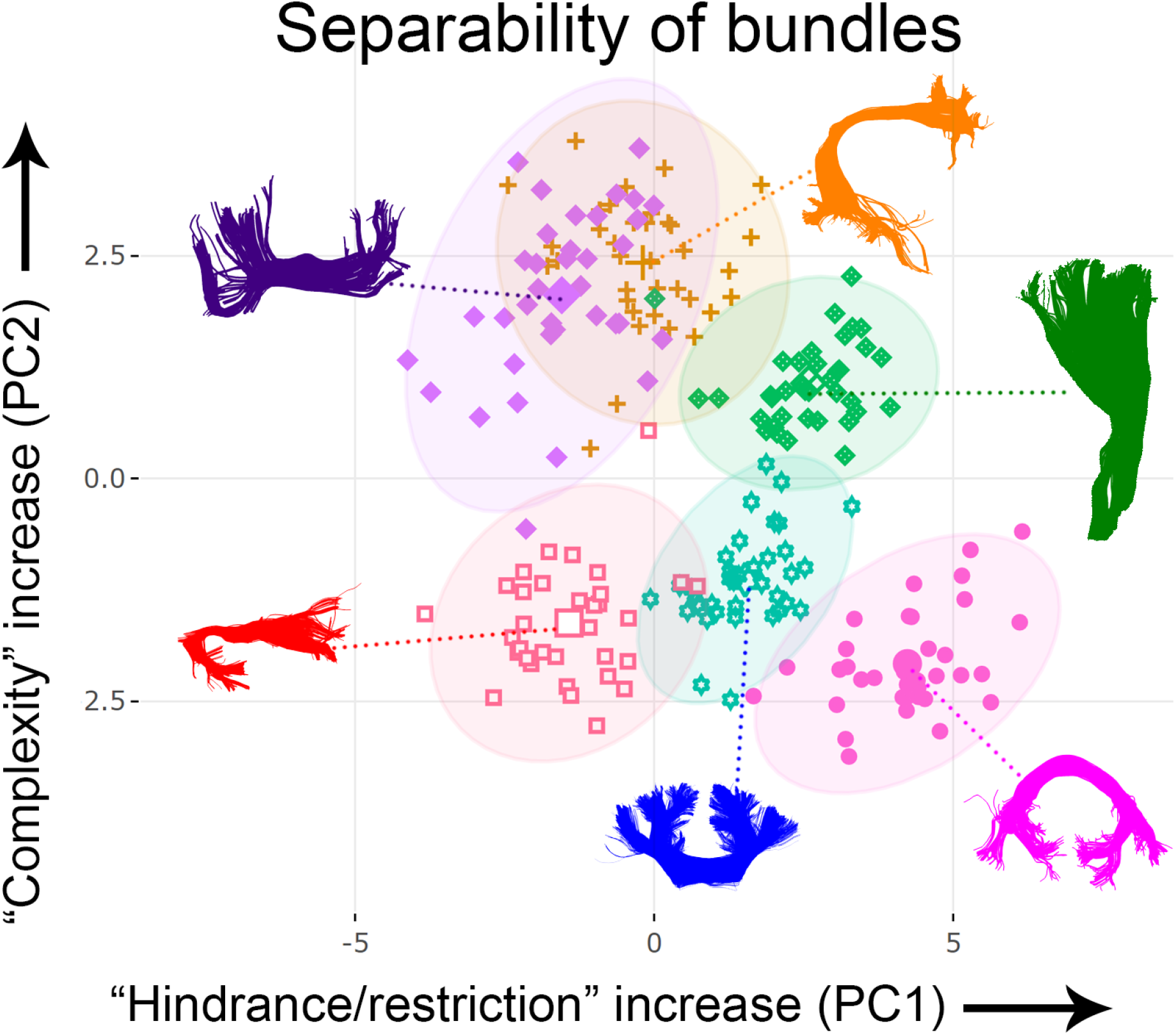
Bundle clustering based on PC1 and PC2. The horizontal axis shows increasing restriction or hindrance perpendicular to the main axis of the bundles. On the right-most part of this axis are located densely-packed bundles such as the CST (green), genu (blue) and splenium (pink). The vertical axis represents the complexity degree of bundles (based on NuFO). On top of this axis are the arcuate fasciculus (orange) and superior longitudinal fasciculus (purple), two pathways which are known to have many crossing regions. Each point represents one subject. Concentration ellipsoids cover 95% confidence around the mean.

## Virtual dissection scheme

The following details a brief description of the virtual dissection plan of each white matter bundle reported in the present study. All bundles were interactively extracted separately, in native space, repeating the method for each subject and hemisphere. Dissection were performed using FiberNavigator (Chamberland et al., 2014) by co-author KD, under the supervision of tractography specialists MC and DKJ.

### Anterior commissure (AC)

The AC is a thin white matter bundle which forms a direct connection between the temporal lobes (Wakana et al., 2007). It is located inferior to the most anterior part of the fornix and was captured by placing ROIs on the lateral branches in the ventral temporal lobe (Catani and De Schotten, 2008).

### Arcuate Fasciculus (AF)

To extract the AF, the first ROI was placed laterally to the corona radiata to capture the frontal-partial fibres. Then, a second ROI was placed posterolateral to the sylvian fissure to capture streamlines extending into the temporal lobe (Catani and De Schotten, 2008). An exclusion ROI was positioned below the curved portion of the bundle the remove spurious connections.

### Mid-body of the Corpus Callosum (CC)

Two ROIs were placed ventral to the location of the cingulum and medial to the lateral ventricles (one in each hemisphere) (Catani and De Schotten, 2008). Exclusion ROIs were used to exclude the Genu and Splenium (i.e., the anterior and posterior sections of the corpus callosum, respectively).

### Corticospinal Tract (CST)

A first ROI was placed at the level of the cerebral peduncle in the axial plane. A second ROI was placed over the central sulcus and capture fibres extending to the primary motor cortex (Chenot et al., 2018).

### Inferior Longitudinal Fasciculus (ILF)

The ILF is an association pathway forming a direct connection between the occipital and temporal lobe (Wakana et al., 2007). A first ROI was placed laterally, in the anterior temporal pole and a second ROI was placed in the posterior temporal lobe (Catani and De Schotten, 2008).

### Optic Radiation (OR)

To capture this bundle, a ROI was placed in the occipital lobe to encompass V1 and V2. A second ROI was planced anterolaterally to the geniculate body (Yamamoto et al., 2005; Chamberland et al., 2017).

### Superior Longitudinal Fasciculus (SLF)

For the SLF, a first ROI was placed on the superolateral aspect of the cingulum. Then a second ROI was placed over the supramarginal gyrus (Kamali et al., 2014).

### Uncinate Fasciculus (UF)

The UF connects the orbito-frontal cortex to the anterior temporal lobe. A first ROI was placed in the anterior part of the temporal lobe. A second ROI was positioned in the inferior medial frontal cortex (Catani and De Schotten, 2008).

### Frontal Aslant Tract (FAT)

To extract this bundle, a selection ROI was placed over the superior frontal gyrus and a second one was placed middle of the inferior frontal gyrus (Catani et al., 2013).

### Cingulum (Cg)

The cingulum is a medial association fibre that is located over the corpus callosum. To extract this bundle, a first ROI was placed immediately postero-superior to the Genu. A second ROI was positioned in the posterior cingulate cortex (Catani and De Schotten, 2008).

### Genu

The genu is a commissural fibre that is the most anterior section of the corpus callosum. To extract this bundle, two ROIs were placed anterolateral to the most rostral portion of the corpus callosum in each hemisphere. This approach was used to capture the anteriorly arching fibres of the genu (Catani and De Schotten, 2008). An exclusion ROI was used to exclude streamlines extending posteriorly to the Genu, which make up the body of the corpus callosum.

### Inferior Fronto-Occipital Fasciculus (iFOF)

The iFOF connects the orbito-frontal cortex and the ventral occipital lobe. A first ROI was placed covering the anterior floor of the external capsule, and a second ROI was placed in the inferior part of the occipital lobe (Catani and De Schotten, 2008).

### Splenium

The splenium is the most posterior section of the corpus callosum, which joins the temporal and occipital lobes of the two hemispheres. Two ROIs were placed posterolateral to the most caudal section on the corpus callosum in the left and right hemisphere(Catani and De Schotten, 2008). An exclusion ROI was used to remove streamlines extending anteriorly to the splenium.

